# Coordinated alternative splicing decisions via stepwise exon definition

**DOI:** 10.1101/2024.12.18.629257

**Authors:** Panajot Kristofori, Simon Braun, Anke Busch, Kathi Zarnack, Julian König, Stefan Legewie, Congxin Li

**Affiliations:** Department of Systems Biology, Institute for Biomedical Genetics (IBMG), University of Stuttgart, 70569 Stuttgart, Germany; Stuttgart Research Center for Systems Biology (SRCSB), University of Stuttgart, 70569 Stuttgart, Germany; Institute of Molecular Biology (IMB), Ackermannweg 4, 55128 Mainz, Germany; Buchmann Institute for Molecular Life Sciences (BMLS), Goethe University Frankfurt, Max-von-Laue-Str. 15, 60438 Frankfurt, Germany; Theodor-Boveri-Institut für Biowissenschaften, Lehrstuhl für Bioinformatik, Biozentrum, Julius-Maximilians-Universität Würzburg, Am Hubland, 97074 Würzburg; Theodor-Boveri-Institut für Biowissenschaften, Lehrstuhl für Biochemie und RNA-Biologie, Biozentrum, Julius-Maximilians-Universität Würzburg, Am Hubland, 97074 Würzburg

## Abstract

Alternative splicing of pre-mRNA is a fundamental step in human gene regulation and aberrant slic-ing is tightly linked to diseases including cancer. Various splicing events, such as alternative exon (AE) choice and intron retention (IR), are controlled by a common molecular machinery, the spliceosome. However, it remains elusive how the regulation of spliceosome activity coordinately affects these splicing decisions. Here, we analyze a large-scale mutagenesis screen and genome-wide RNA sequencing data to show that IR and AE choice are tightly coupled, as IR products primarily accumulate in alternative exons showing intermediate inclusion levels. Using data-driven mathematical modeling, we reveal that multistep exon recognition by the spliceosome explains the observed AE-IR dependency for cis-acting sequence mutations and upon knockdown of trans-acting RNA-binding proteins. Furthermore, we show that multistep exon definition is frequently perturbed in cancer cells, which leads to the coordinated deregulation of intron retention and AE choice. In conclusion, we showed that the spliceosome coordinates complex splicing decisions via stepwise exon definition, which potentially facilitates the search for common molecular mechanisms for mis-splicing in cancer.

## Introduction

Alternative splicing (AS) of pre-mRNA amplifies the diversity of the transcriptome and proteome in most eukaryotes, allowing for robust and versatile gene expression involved in cell physiology and development. In humans, over 90% of the gene transcripts are alternatively spliced, yielding an average of 3.4 isoforms per gene (1–3), and in some cases, thousands of isoforms can emerge (e.g., neurexins). The diverse isoforms are produced via a relatively small number of modular AS events, such as inclusion/exclusion of the alternative exon (AE), usage of alternative 5’ or 3’ splice sites (SSs), intron retention (IR). Among these AS events, intron retention was largely overlooked in the mammalian systems, as it was generally thought as a mis-splicing incidence that triggers the clearance of the IR isoforms via nonsense mediated decay (NMD) and thus provides no physiological significance. However, recent experimental and computational advances have revealed that IR actively renders regulatory roles in many biological functions, e.g., facilitating prompt cell response to stress (4, 5) as well as coordinating gene expression program in cell differentiation (6, 7). At the transcriptome-wide level, IR was found to be more prevalent than previously assumed and it was the most sensitive AS event to perturbations in splicing regulators (8). Particularly, IR is the most commonly upregulated type of alternative splicing upregulated in cancer (9). It is thus of great interest to understand the regulatory rules of IR and its relationship with other splicing events such as AE usage.

Splicing is carried out by the spliceosome, a large ribonucleoprotein complex that undergoes progressive assembly and remodeling on a transcripts. Initially, spliceosomal U1 and U2 small nuclear ribonucleoproteins (snRNPs) bind to the 5’SS and the branch point (BP) upstream to the 3’SS, respectively. Further recruitment of the U4/U6.U5 tri-snRNP leads to the formation of a cross-intron complex (B complex) establishing 5’ -3’SS communication. The B complex then undergoes a series of energy-dependent remodeling and becomes active for subsequent intron-removal catalysis. This type of splice-site recognition and pairing via forming cross-intron complex is referred to as intron definition (10). Another spliceosome assembly path is mediated by the initial formation of a cross-exon complex. This process, termed exon definition, is preferentially used in human cells, especially for transcripts short exons and long introns w. After cross-exon formation of a B-like spliceosome complex, extensive ATP-dependent remodeling and structural rearrangements are required for the conversion to cross-intron spliceosome that is essential for splicing catalysis (11–15). A common feature of both working modes is the ordered multi-phase maturation of the spliceosome driven by energy expenditure, implying the existence of several potential rate-limiting steps. Interestingly, depletion of the spliceosome components and regulators involved in early splice-site recognition steps (e.g., U1 and U2 factors) mainly affects AE usage, whereas perturbations in later steps, such as splice-site communication and spliceosome remodeling, preferentially change IR level (16). Thus, a mechanistic understanding of spliceosome regulation is essential for studying AS decisions.

Splicing decisions are tightly controlled by *cis*-and *trans*-regulators. A compendium of sequence motifs, to which core spliceosome components and specific RNA binding proteins (RBPs) are recruited, has been identified as enhancers and silencers for various splicing events. Very often these elements function in combination in a context-dependent manner, therefore posing a challenge to quantitatively predict their effects on the splicing outcomes. Computational algorithms have been actively developed to integrate these sequence features into predictive models employing a wide range of machine learning techniques and deep-learning neural network architectures (17). Trained by ever-accumulating genomic sequencing data, these models were optimized to quantitatively predict the splicing decisions directly from primary sequences in a tissue-specific manner sequence (18–23). However, these models mainly focus on binary decisions, e.g., included or skipped AE, removed or retained intron, and are thus limited to describing complex splicing patterns with multiple distinct decisions.

Indeed, over one third of the splicing events in humans show complex splicing patterns combining multiple AS decisions at the same splice junction (24, 25). Large-scale mutagenesis experiments showed that even a three-exon minigene can produce multiple isoforms by combining different splicing decisions already in WT and that mutations may activate cryptic splice sites further enriching the splicing choices (26, 27). Thus, a single metric, such as PSI (percent spliced-in) for AE, is not sufficient to quantify the complex splicing decisions. To address this issue, kinetic models have been developed to describe the generation of multiple isoforms from the same transcript combining several splicing events (26–30). These models allow for analyzing the regulation of not only binary but multiple splicing outcomes. However, a quantitative description of how intron retention coordinates with other splicing decisions is still missing for specific genes and transcriptome wide.

In this work, we show that the intron retention and AE choice are coordinately regulated in a non-monotonic way, where intron retention accumulates strongly at intermediate AE inclusion level. This regulatory dependency was observed in a RON minigene subjected to large-scale mutagenesis as well as in genome-wide data from over 20 human tissues. We proposed a mechanistic model driven by the RON mutagenesis data and found that the multi-step AE definition underlies the coordinated splicing regulation. We further show that the strength of the AE-IR dependency is most sensitive to RBPs differentially modulating the AE definition steps, reflecting the regulatory roles of RBPs in different stages of spliceosome assembly and maturation. At the transcriptome-wide level, the differential modulation of AE definition steps also accounts for the shift in AE-IR dependency in many cancer types, implying heterogeneous perturbations on exon recognition by malignant cell states. Taken together, we provided evidence of coordination between two important splicing events and showed its relevance to analyzing alternative splicing regulation in perturbed cellular contexts such as cancer.

## Results

### Alternative exon and intron retention events are coordinately regulated

To investigate the interplay between the modulations of multiple splicing events, we made use of our previously published data from a high-throughput mutagenesis screen of minigenes harboring RON exons 10-12 (26, 30). The pre-mRNA of RON minigene is alternatively spliced into five major isoforms – AE inclusion, AE skipping, first intron retention, second intron retention and full intron retention, and we had quantified the impact of 1800 point mutations on the abundance of these isoforms (26, 30). Here we focus on the interdependency of AE choice and intron retention characterize the link between the two events using two metrics for individual mutations (Figure 1A). First, the percent spliced in (PSI) of the alternative exon, i.e., PSI = *f*_inclusion_ / (*f*_inclusion_ + *f*_skipping_) with *f* repre-senting isoform frequency, quantifies the specificity of the binary alternative exon choice ranging from total skipping (PSI = 0%) to full inclusion (PSI = 100%). Second, the efficiency of intron removal, defined as efficiency = *f*_inclusion_ + *f*_skipping_, measures the overall occurrence of active splicing and it is complementary to the total level of intron retention, i.e., efficiency = *f*_inclusion_ + *f*_skipping_ = 100% -(*f*_IR1_ + *f*_IR2_ + *f*_fullIR_).

**Figure 1.**
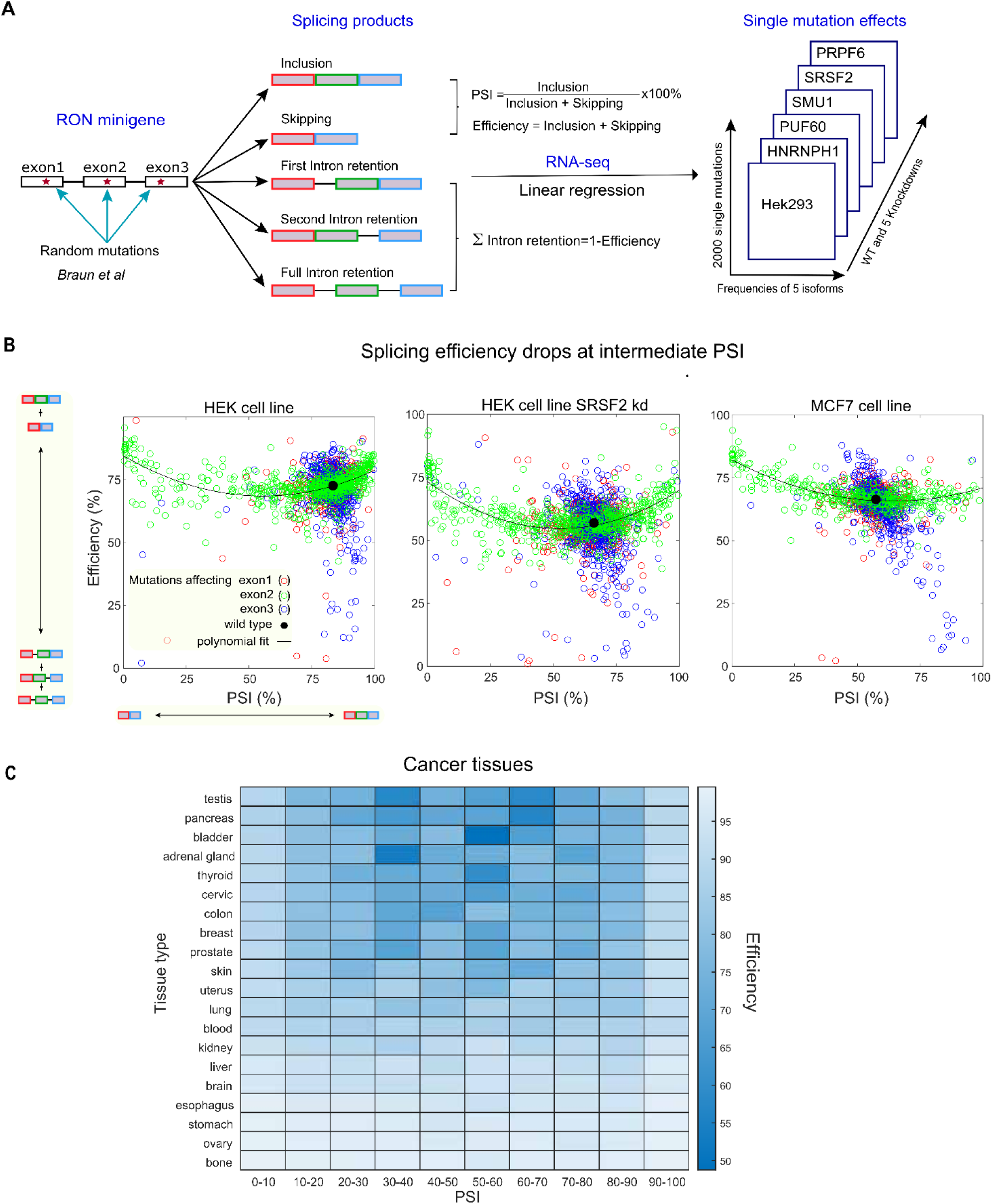
Splicing efficiency drops at intermediate alternative exon inclusion level. **(A)** Published mutagenesis data used in this study. Braun et al. performed linear regression analysis on the random mutagenesis data of the RON minigene comprising exons 10-12 (here designated as exons 1-3) to quantify the single-mutation effects on abundance of five main splice isoforms, AE inclusion, AE skipping, first, second and full intron retention. Two metrics were used to characterize splicing outcomes for each point mutation: PSI quantifies the binary choice of AE usage, and the splicing efficiency quantifies the summed frequencies of AE inclusion and skipping and is complementary to the sum of all intron retention isoforms. The analyzed data from Hek293 cells is a matrix containing the frequencies of five splice isoforms for around 1800 single point mutations, for control and five RBP knockdowns, namely HNRNPh1, PUF60, SMU1, SRSF2 and PRPF6 (the last 4 being non-published data; see Methods for details). **(B)** Splicing efficiency drops at intermediate PSI for various experimental conditions in RON minigene. The plots depict PSI and efficiency across a spectrum of 1800 mutations. Each dot represents one single point mutation: red and blue indicate mutations in outer exons, respectively, and green represents mutations in the AE. The black dot represents WT RON minigene without mutations. The black line represents a polynomial fit for the AE mutations. Left: HEK293 cell line; middle: SRSF2 knockdown in HEK293 cells; right: MCF7 cells. **(C)** Splicing efficiency drops transcriptome-wide in cancers. Heatmap shows the median efficiency at different bins of PSIs in 20 cancer types, where the data were acquired from MAJIQLOPEDIA. The cancer datasets were from TCGA, TARGET, BEAT-AML and CBTN different healthy tissue sub-brain regions.

As reported previously (30), point mutations in the outer constitutive exons lead to strong accumulation of intron retention products (low splicing efficiency), while having little effect on PSI (Fig. 1B, blue and red dots). Point mutations located to the AE strongly affected the PSI, but also affected the splicing efficiency (Fig. 1B, green dots). Specifically, we observed a drop to ∼ 65% efficiency for mutations establishing intermediate PSI levels when compared to the much higher splicing efficiencies (90%) observed for mutations inducing extreme PSIs (0 or 100%). Hence, while intron retention can be independently regulated, the modulation of the alternative exon is accompanied by the accumulation of intron retention products.

To further test whether this splicing efficiency drop is a robust phenomenon, we generated and analyzed RON mutagenesis data in different cellular contexts, including siRNA-mediated knockdown of several RBPs (HNRNPH1, PUF60, SMU1, SRSF2 and PRPF6; Figure 1A) and using another cell type (MCF7 cells; Methods). The non-monotonic dependency of splicing efficiency on PSI was observed in all data sets (Figure 1B and Supplementary Figure S1). Interestingly, the efficiency drop is quantitatively different across conditions, showing large (SRSF2 and SMU1), intermediate (PRPF6) or no change (HNRNPH1 and PUF60) compared to that in WT HEK293 cells (Figure 1B left panel; Supplementary Figure S1). Thus, the PSI-dependent splicing efficiency drop is an intrinsic property of AE regulation that is subject to regulation by trans-acting factors.

To investigate whether this coordinated splicing regulation is a general feature observed for genes besides RON, we extracted and analyzed exon junctions showing both AE and intron retention events from published genome-wide RNA-seq data in 20 human tissues assembled in MAJIQLOPEDIA (31) (Methods; Supplementary Table S1). For quantification, we calculated the average splicing efficiency of all exons within 10 PSI bins ranging from near-complete skipping (PSI_0-10%) to near-complete inclusion (90-100%) in normal and cancer tissues. In line with the observations for the RON minigene, the average splicing efficiency drops at intermediate PSIs in 15 out of 20 tissues (Figure 1C and Supplementary Figure S1). Interestingly, we also noticed that the IR declines in 5 out of 20 cancer types (Figure 1C), indicating that cancer disturbs IR decision in a tissue-dependent manner.

Taken together, we found that the alternative exon and intron retention events are in general coordinately regulated, where balanced alternative exon inclusion and skipping leads to strong accumulation of intron retention products. This phenomenon was commonly observed not only in the RON minigene reporter but also genome wide in various genetic and cellular contexts. It thus raises questions about the molecular mechanism underlying this behavior.

### Two-step exon definition explains interdependence between alternative splicing events

To understand the mechanism of the interplay between alternative splicing events, we develop quantitative mechanistic models and statistically test their abilities to faithfully capture the data of point-mutation effects on RON splicing. Following up our previous work (30), we studied an exon-definition model of RON splicing, in which each of the three exons in the minigene is defined via cooperative binding of spliceosomal U1 and U2 and forming cross-exon complex (Figure 2A). The U1-U2 assembly across an exon is described as a one-step reversible association-dissociation process (Figure 2A, one-step model in the green bubble). Assuming independent definition of the three exons, we considered 8 binding states in total and assumed that these dictate splicing outcomes. Specifically, pre-mRNA with all three exons defined gives rise to AE inclusion isoform, and AE is skipped when only the two outer exons are defined. An undefined first or last exon, with the other two exons defined, leads to first or second intron retention, respectively. All binding states can be converted into full intron retention isoform, reflecting that unproductive splicing or nuclear export of a transcript leads to terminally retained introns (detailed model scheme see Supplementary Figure S2). The binding affinities to the three exons shape the distribution of the 8 spliceosome binding states and thus the concentrations of the splicing products. As the spliceosome binding is regulated by sequence context and RBPs, the model can be used to study the effects of genetic perturbations, e.g., mutations and RBP depletion, on the splicing outcomes.

**Figure 2.**
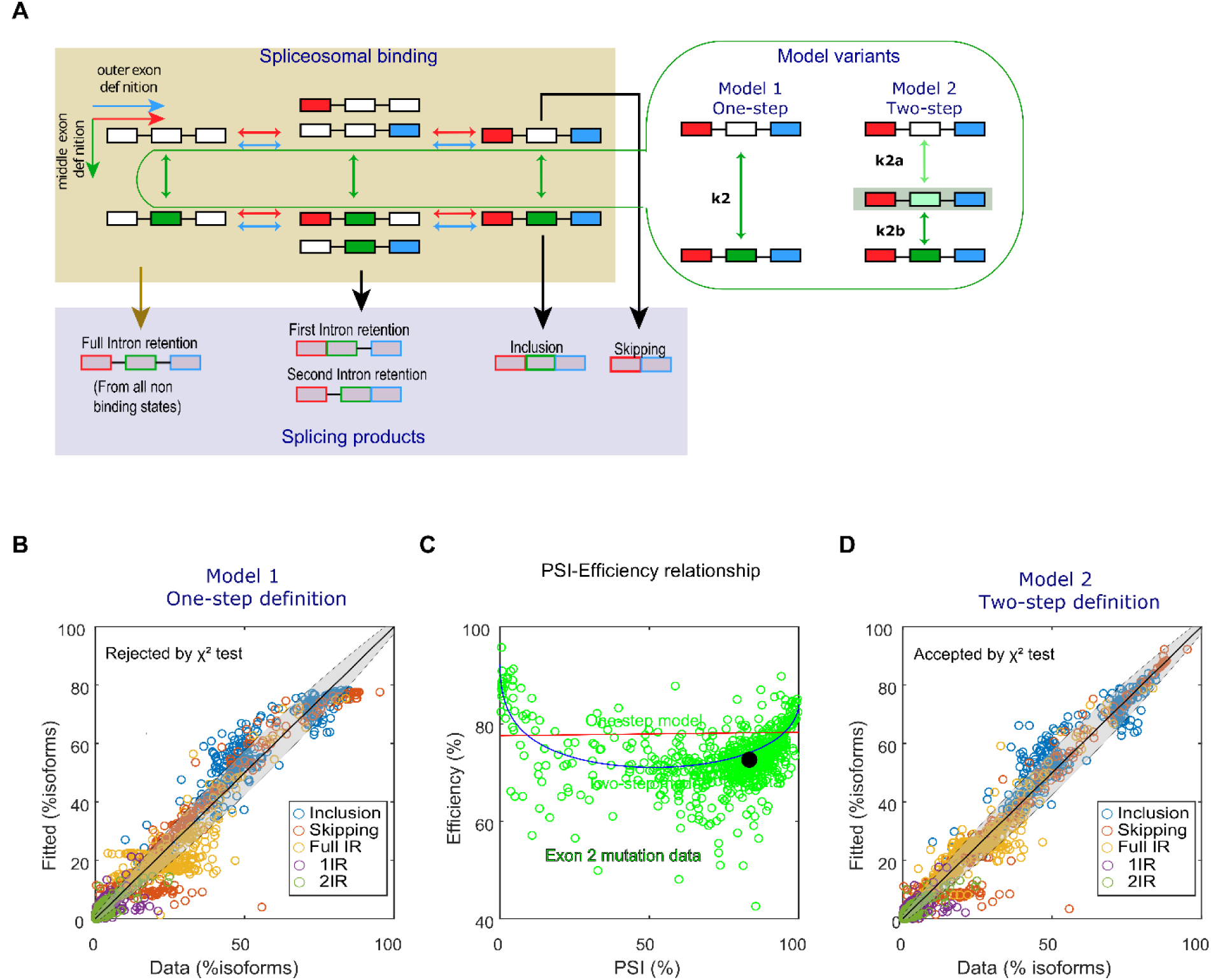
The two-step exon definition model explains splicing efficiency drop at intermediate PSI level. **(A)** Schemes of one-step and two-step AE definition models. The illustration inside the yellow shading depicts the independent definition of the three exons in the minigene by spliceosome binding following the exon definition mechanism. Empty and filled boxes represent undefined and defined exons, respectively. The two models differ only in the AE definition mechanism, with one or two rate-limiting steps (green bubble and arrows), while the definition of the two outer exons follow the same one-step process (blue and red arrows). Exon definition states determine splicing outcomes (blue shading): Pre-mRNAs with all three exons defined give rise to AE inclusion isoform, and those with only AE undefined skip AE in the final product. If only the first or the last exon is left undefined, first or second intron retention isoforms will be generated, respectively. All spliceosome binding states can be converted to full intron retention in both models. **(B)** One-step model fails to fully describe the single-mutation effects on RON splicing. The one-step model was fitted to the frequencies of all five RON isoforms for 1800 single mutations. The model was rejected by the *χ*^2^ goodness of fit test. Black lines: the diagonal line representing perfect data-model agreement; shade: isoform variability predicted by the error model. **(C)** Coordinated AE-IR regulation described by the two-step and not the one-step model. The data in green circles represent the mutations in the AE of the RON minigene and the lines were from the best fits of the one-step (red) and two-step (blue) model, respectively. Black dot: WT. **(D)** Two-step model faithfully captures the single-mutation data. The two-step model was fitted to the frequencies of all five RON isoforms for 1800 single mutations. The model was accepted by the *χ*^2^ goodness of fit test. Same representation as in B.

To test whether the one-step definition model can explain the point-mutation data of RON minigene in HEK293 cells, we translated the model scheme into ordinary differential equations (ODEs) and fit the steady-state solution for the frequencies of the five splice isoforms to the corresponding data of RON minigene (standard *χ*^2^ fit), with the constraint that each mutation only affects the spliceosome association rate of its nearest exon (Methods). For instance, mutations in or around exon 2 (± 40 nt) are assumed to solely affect the exon definition rate *k*_2_. While the isoform frequencies of the best fit seemed to follow the tendency of the measured ones, the one-step definition model was rejected by the *χ*^2^ goodness-of-fit and thus failed to fully explain the data (Figure 2B; Methods). Indeed, the model inherently features the independent regulation of alternative exon and intron retention (Figure 2C, the red flat line), so it qualitatively fails to describe the interplay between the two splicing events even with optimized parameters.

To overcome this limitation, we extended the exon definition model. Recent studies reported that the spliceosome conformation undergoes extensive energy-dependent remodeling before splicing catalysis (12–15, 32), which may lead to multiple rate-limiting steps during exon definition. To incorporate this, we extended the model assuming that the definition of the alternative exon requires two consecutive steps with different rates (Figure 2A, two-step model in the green bubble). The intermediate state between the two steps reflects immature or uncommitted spliceosome assembly that potentially hinders subsequent splicing catalysis and thus only yields the unspliced isoform, i.e., full intron retention. We fitted this model with the same point-mutation data assuming that U1 and/or U2 act in both steps of AE definition and thus a mutation in or near exon 2 affects the rates of both steps (Methods). The two-step definition model is accepted by the *χ*^2^ test and thus quantitatively captures the data, (Figure 2C). Furthermore, the best fit qualitatively reproduced the splicing efficiency drop at intermediate AE inclusion level (Figure 2B, blue arc).

In conclusion, by iterative data-driven modeling, we showed that the two-step exon definition model reproduces the coordinated regulation between AE and IR events, suggesting that multiple rate-limiting steps occur in the regulation of spliceosome decisions.

### Two-step exon recognition combines isoform switching with high splicing efficiency

While the two-step exon recognition well explained the AE-IR dependency (Figure 2), it remained unclear whether and how this mechanism provides any functional significance. To investigate the potential benefit of the two-step regulation for alternative splicing, we performed numerical simulations in which we induced a shift from skipping to inclusion by varying the AE recognition parameter(s) (from the best fit values). In these simulations, we compare one-and two-step mechanisms with respect to the PSI response curve. Even though altered recognition of the AE induces a complete shift from skipping to inclusion (PSI from 0 to 100%) in both models, PSI responses more sharply in the two-step model (Hill coefficient *h* = 1.8) than in the one-step model (*h* = 1.1). Thus, in the two-step model a relatively small change in AE recognition due to mutations or other regulatory mechanisms is sufficient to effectively change splicing outcomes. Specifically, a 11-fold change in AE recognition switches the PSI from 10% to 90% (Figure 3A), whereas one-step model requires much larger change in AE recognition (50 fold) to achieve the same PSI switching. Therefore, the two-step AE definition enables a more efficient splicing switch.

**Figure 3.**
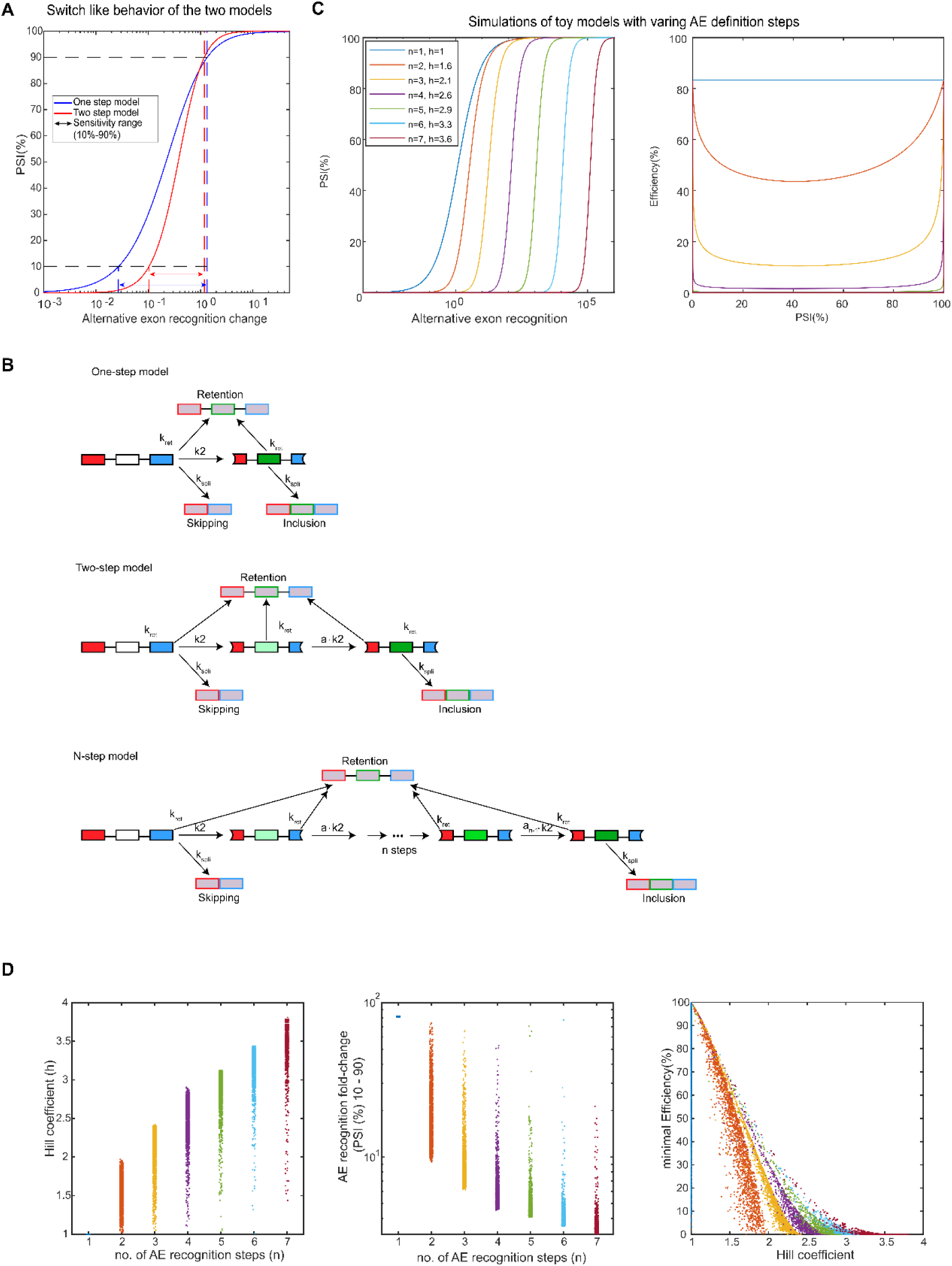
Two-step model balances a trade-off between sensitive switching and high efficiency in splicing regulation. **(A)** Response of PSI to changes in the AE definition rate simulated by the one-step (blue curve) and two-step (red curve) models. The simulation was performed by varying only the AE definition rates (*k*_2_ in the one-step model; proportional change in *k*_2a_ and *k*_2b_ in the two-step model) starting from the best-fit parameters. To switch the AE inclusion level (PSI) from 10% to 90% (horizontal dashed lines), an 50-(blue dashed lines) and 11 (red dashed lines) fold-change in the AE definition parameters is required in the one-step and two-step models, respectively. **(B)** Toy models describing one-step, two-step and *n*-step AE recognition. The two outer exons were assumed to be strongly recognized (i.e., constitutive exons) and only AE recognition was modelled with different numbers of binding steps (from 1 to 7). For multi-step models (*n* ≥ 2), the recognition rates were assumed to change stepwise by a same fold-change factor (*a*) between consecutive steps, i.e., rate of the *i*th step = *k*_2_*a*^*i*−1^, and *a*<<1 was assumed to reflect efficient proofreading. The fully defined and undefined AE states give rise to AE inclusion and skipping isoforms, respectively, with a common splicing rate *k*_spli_. Another alternative splicing decision for all recognition states is the retention of introns in the final product (with the rate *k*_ret_). **(C)** Multi-step AE definition strengthens the switch-like AE regulation at the cost of reducing splicing efficiency. Left panel: dose-response curves of PSI as a function of AE recognition strength (*k*_2_) were simulated for a range of intermediate AE recognition steps (from 1 to 7). The one-step model showed the weakest switch-like regulation (Hill coefficient *h* = 1), whereas multi-step models allowed for a sharper switch (Hill coefficient *h* ranging from 1.6 to 3.8) with increasing number of steps (from 2 to 7). Right panel: However, the same simulations showed that the splicing efficiency stronger drops if the number of intermediate steps is larger than two, which reflects strong accumulation of unproductive splicing products. **(D)** Systematic parameter sampling confirms the trade-off between switch-like AE regulation and splicing efficiency for a large range of parameter values. All parameters of the toy models were randomly sampled (see Methods), and for each simulation the Hill coefficient and the minimal splicing efficiency for varying k_2_ were calculated as in (C). In line with the results in (C), the switch-like regulation (quantified by the Hill coefficient) increases and the sensitive switching interval of AE recognition strength (causing 10% to 90% change in PSI) decreases with the number of intermediate AE recognition steps (left and middle panels, respectively). Furthermore, the trade-off between switch-like AE modulation and splicing efficiency still holds true (right panel). Same color code as in (C).

The two-step model explains the AE-IR dependency and potentially facilitates switch-like splicing regulation, and it raises a question whether introducing even more steps in the AE definition would further improve the model fit and PSI switching. To address this question, we focused on the core module of AE definition and formulated toy model variants with different complexity by varying the number of intermediate recognition steps (one to seven) during the transition from an undefined to a fully defined AE (Figure 3B). In all model variants, the first and the last species in the chain give rise to skipping and inclusion products, respectively, whereas productive splicing is not possible from the partially defined AE intermediates. By simulations using the best-fit AE definition parameters, we observed almost the same Hill coefficients in the one-and two-step toy models as in the corresponding full models, confirming that the toy models faithfully characterize AE switching from skipping to inclusion.

Next, we simulated the model variants with three to seven AE definition steps (Methods). The corresponding Hill coefficients range from 2.1 to 3.6, implying that a 6 to 3-fold change in AE recognition is required to switch PSI from 10 to 90%, respectively (Figure 3C left). Compared to the difference in switch-like behavior between the one-and two-step models, these values of the three to seven-step models are close to that of the two-step model. However, increasing the number of AE recognition steps comes at a cost for splicing efficiency, as the IR level drastically increases with the number of molecular steps (Figure 3C right). Based on this analysis, we conclude that the gain in switch-like splicing behavior is limited for a larger number of AE recognition steps.

To test this result in a more general context, we randomly sampled the kinetic parameters in all toy models and quantified the PSI response for each parameter set to estimate the range of Hill coefficients that can be attained in each variant (Figure 3D; Methods). Indeed, the gain in isoform switching realized by one additional molecular step continuously decays with the increasing number of molecular steps. Hence, increasing the number of exon definition steps beyond two has a limited effect on switch-like PSI modulation and therefore the potency of splicing regulation for a given fold-change in AE recognition. Moreover, for models with more than two molecular steps, we observed a very strong accumulation of unspliced precursors/products at intermediate PSI. Thus, a high number of intermediate steps leads to problems with splicing execution, slowing down the reaction and/or leading to the accumulation of products with terminal intron retention.

Taken together, we show here that two-step AE recognition may meet a trade-off, combining sensitive isoform switching and high splicing efficiency. Even though a larger number of AE definition steps may improve switch-like behavior and splicing fidelity (rejection of suboptimal substrates), this comes at the cost of drastically reduced splicing efficiency. We therefore suggest that the number of steps in AE recognition is limited to ensure potent alternative splicing regulation at high splicing efficiency.

### Identification of RBP regulatory roles by modeling coordinated AE and IR events

Next, we sought to understand how the splicing-event interdependency is regulated and what could be learned from it about molecular mechanisms of splicing regulation. To reflect our data, in which RBP knockdown effects are assessed across the whole spectrum of mutations (Supplementary Figure S1), we simulated how the efficiency-PSI dependency shifts when a RBP controls molecular steps in the best-fit full two-step exon-definition model (Figure 2D and Methods). RBPs exclusively regulating the AE showed two distinct modulation modes: If a RBP controls both AE definition steps concertedly, i.e., altering individual rates but maintaining their ratio, a perturbation of the RBP abundance will leave the efficiency-PSI curve unchanged and only shift the starting point representing the unmutated RON minigene upon RBP KD (Figure 4A). Alternatively, a RBP differentially modulating the two AE definition steps, e.g., modulating only one step, controls not only the position of the WT minigene but also the shape of the efficiency-PSI dependency curve, while maintaining the efficiencies at extreme PSIs (Figure 4B). In contrast, knockdown of a RBP regulating the definition of the outer (constitutive) exons causes a general shift of all efficiency levels coupled with shape change of efficiency-PSI curve (Figure 4C). Interestingly, we also observed similar patterns in the RPB KD data: first, the knockdowns of PRPF6 and PUF60 change only the WT position, but leave the PSI efficiency curve unaffected as expected for concerted AE modulation (Figure 4D; related to Figure 4A); second, the effects of SRSF2 and SMU1 knockdowns suggest that these are general splicing regulators, which induce combined regulation of the outer exons and the AE, as both WT position and the efficiency-PSI dependency are strongly perturbed (i.e., Figure 4E resembles a combination of simulated effects in Figure 4B and C). This implies that quantitative shifts in efficiency-PSI dependency can be used to identify the splicing step(s) a RBP regulates and the directionality of the regulation (i.e., activator or inhibitor).

**Figure 4.**
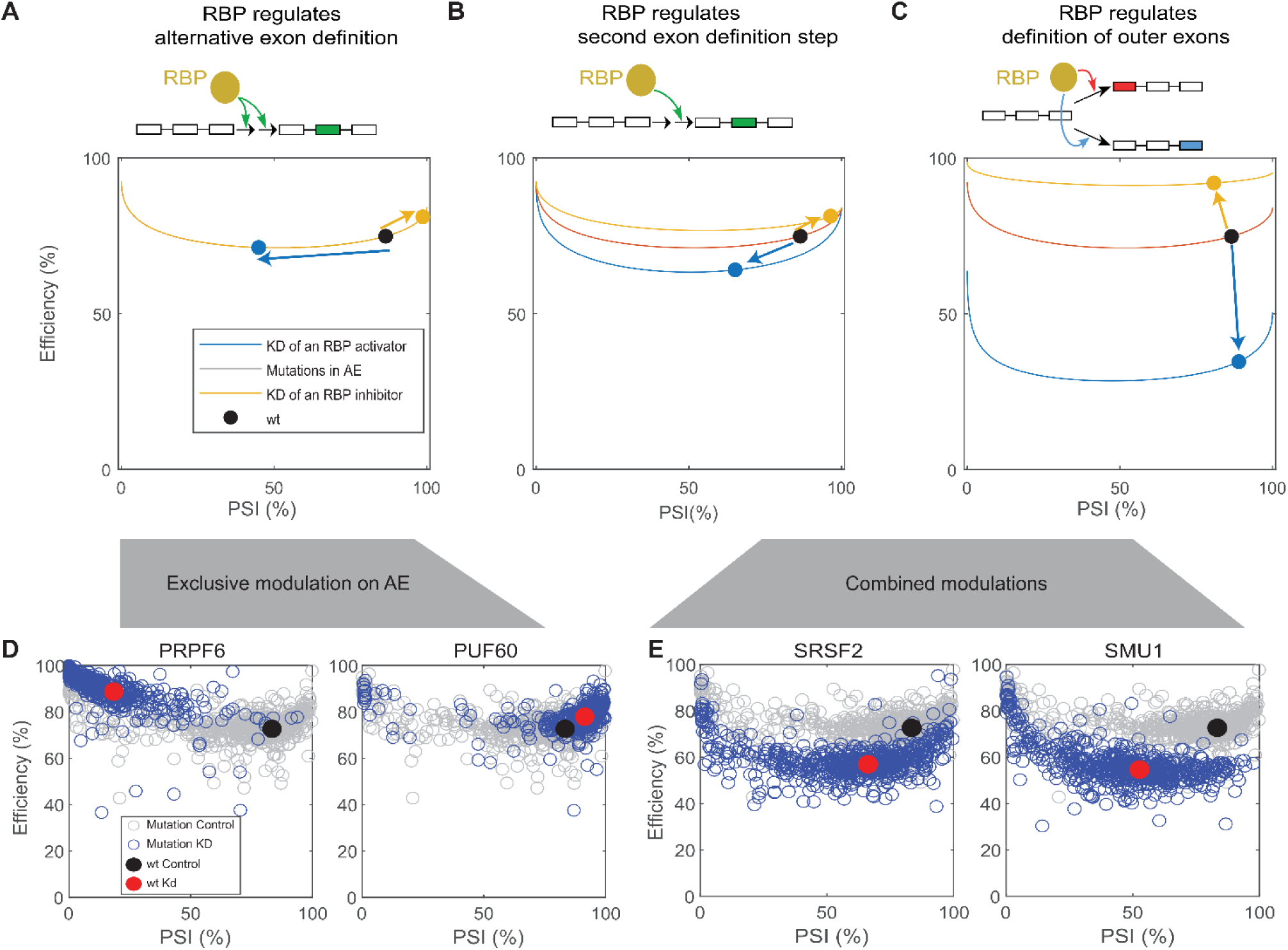
RBP KDs have qualitatively different effects on efficiency-PSI dependency. (**A** – **C**) Qualitative simulations were performed to mimic knockdowns of RBPs regulating different exon-definition steps in the two-step model. The parameters from the best-fit of the RON minigene (Figure 2D) were altered according to the following regulation modes: RBPs can exclusively modulate AE by changing proportionally both AE definition rates (A) or only one of them (B). In addition, RPBs can also affect the definition of the two outer exons (C). The black dot and the red curve represent the efficiency-PSI dependencies in the unmutated WT control and simulated exon 2 mutants, respectively. The blue and yellow curves indicate corresponding shifts simulated for a KD of RBPs acting as an activator (blue) or a repressor (yellow). (**D** and **E**) PSI-efficiency shifts observed in RBP KD data for the RON minigene resemble the simulations in (A – C). (**D**) Compared to scrambled KD control (grey circles and black dot), the PRPF6 and PUF60 KDs (blue circles and red dot) mainly shift the position of unmutated WT minigenes (dots), while leaving the trend of mutated minigenes unaffected (circles). This behavior resembles exclusive AE regulation simulation in (A). (**E**) SRSF2 and SMU1 KDs globally change the efficiency-PSI relation for both unmutated and mutant minigenes, mimicking a combined modulation on both AE and outer exons (combination of simulation scenarios in B and C). Same representation as in (D)

Further inspection of the full data considering isoforms further supported our hypothesis that coordinated shifts of splicing outcomes can be used to distinguish modes of RBP regulation. For example, the depletion of PUF60, PRPF6 or HNRNPH1 or mainly affects inclusion and skipping isoforms (Figure 5A, left panel; Supplementary Figure S3), whereas SMU1 or SRSF2 KD leads to an accumulation of full intron retention at the expense of inclusion isoforms (Figure 5A, right panel; Supplementary Figure S3). To identify the RBPs’ regulatory roles in a more quantitative and systematic way, we built a family of models (15 model variants in total) in which a RBP can modulate all possible combinations of exon-definition steps and confronted all model variants with all isoform data for each RBP KD (Figure 5B). To select the best model(s), we assessed both the goodness-of-fit and predictive power: first, we fitted each model variant to the measured isoform patterns of the WT minigene upon RBP KD (frequencies of five isoforms without mutations); second, we cross-validating the fitted results via prediction of the isoform frequencies of all mutant minigenes upon the same RBP KD (using inferred mutation effects in WT RON minigene in HEK293 cells; see Method for details; Figure 5B). To evaluate the quality of model fitting and prediction, we made use of the *χ*^2^ test statistics, i.e., the ratio between the *χ*^2^ value for model fitting (or prediction) and the critical *χ*^2^ for model rejection test (with corresponding degrees of freedom at *p* = 0.05; Methods). This unbiased systematic model selection identified three regulation modes for the tested RBPs: exclusive AE regulation in concerted (HNRNPH1 and PUF60) and differential (PRPF6) manner, as well as combined outer-exon and differential AE modulation (SMU1 and SRSF2). The fitting and prediction from the best models agreed well with the KD data for all tested RBPs (PUF60 and SMU1 KDs in Figure 5B; others in Supplementary Figure S3). The model also captured the characteristic shifts in efficiency-PSI dependency for distinct regulator types as predicted theoretically (Figure 5C compare with Figure 4A-C). This raises an important point that while the change in PSI informs the directionality of the RBP regulation (Figure 5D, blue dots), further quantification of splicing efficiency (or intron retention) is required to fully distinguish the regulatory modes of different RBPs (Figure 5D, red dots).

**Figure 5.**
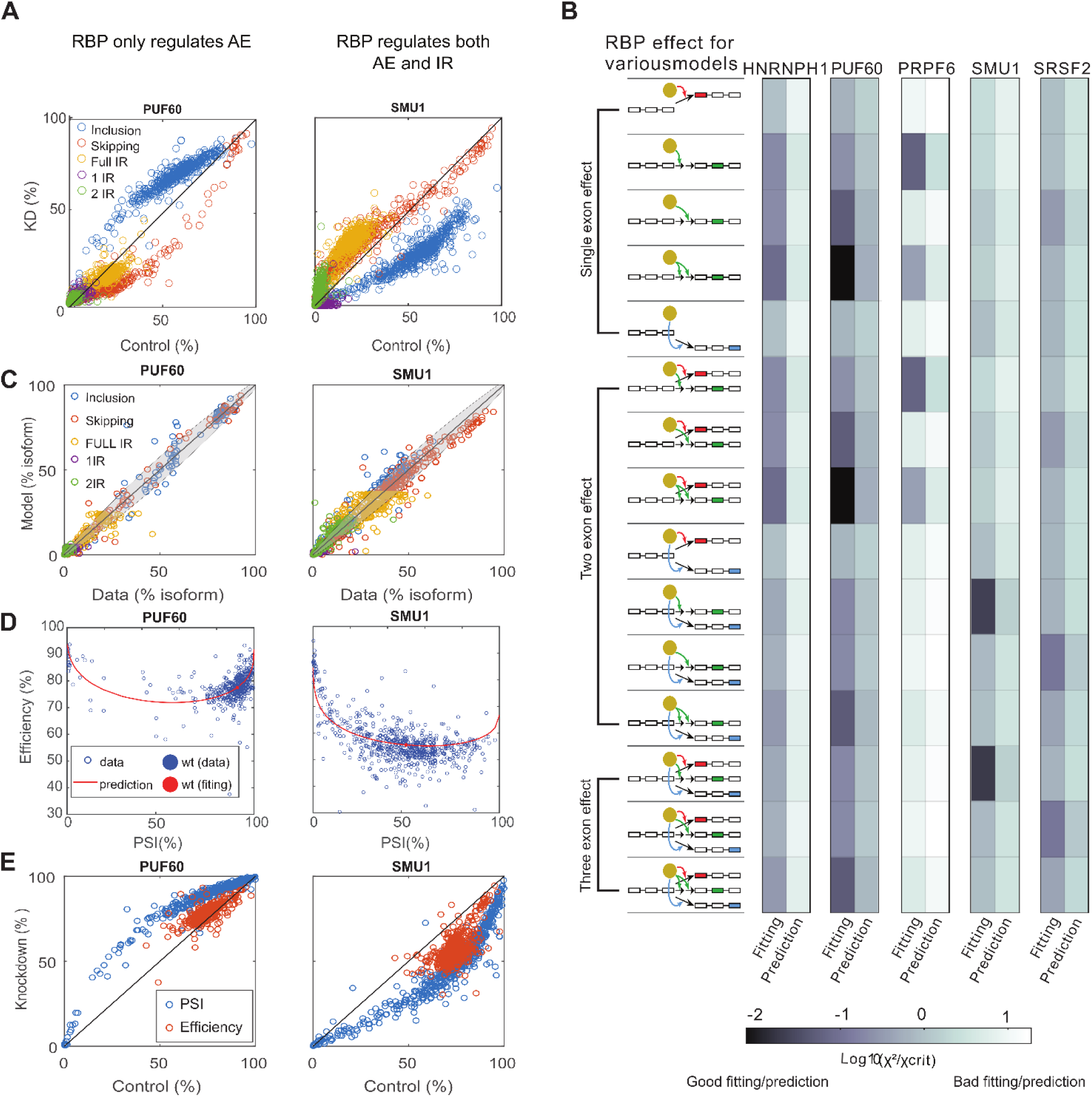
Systematic model selection reveals distinct regulatory roles of RBPs. **(A)** Data of the frequencies of the five RON isoforms shifted upon RBP KDs in the 1800 single mutants (each circle representing a single mutation). PUF60 KD mainly changes the AE inclusion (blue) and skipping (red) isoforms (left panel), whereas SMU1 KD additionally shifts IR isoforms (right panel). **(B)** Computational model selection identifies regulation modes of distinct RBPs. 15 model variants were generated according to all possible combinations of RBP modulation on each exon-definition step, as indicated by the schemes on the left (RBP: yellow dot; RBP-mediated regulation: arrows). For each RBP KD, the unmutated WT minigene was fitted by all 15 model variants, starting from the best-fit model (Figure 2D) and re-fitting only the assumed RBP-regulated parameters. The mutant data (1800 single mutations) was used for cross validation to test the predictive power of each model variant (see Methods for details). Model variants are compared with respect to the fitted and the predicted *χ*^2^ values relative to their corresponding critical values used for *χ*^2^ test (*χ*^2^ ratio for color code). **(C)** Best models combining good fit and high predictive power selected in (B) successfully capture the frequencies of all five splice isoforms in 1800 single mutants and the unmutated minigene for PUF60 (left) and SMU1 (right) KDs shown as examples. Color code: five isoforms. **(D)** Coordinated efficiency-PSI regulation is reproduced by the best-fit models in (C). Blue and red filled circles represent measured and fitted unmutated WT minigene, respectively. Blue open circles and red curves are the measured and model predicted data for single point mutants, respectively. **(E)** PSI (blue) and efficiency (orange) jointly describe the regulatory roles of RBPs when compared between KD scrambled control (x) and RBP-targeted KD (y) for WT and all single mutants (circles). PSI alone can only identify the directionality of a RBP affecting the AE, e.g., PUF60 as a repressor (left) and SMU1 as an activator (right). Additional quantification on efficiency adds another dimension for identifying more detailed modes of regulation (PUF60 KD: no efficiency change; SMU1 KD: efficiency reduction).

Taken together, we found that the dependency of splicing outcomes can be differentially modulated by RBPs acting in distinct exon-definition steps, which provides a basis for quantitatively discriminating the regulatory roles of various RBPs in alternative splicing regulation.

### Two-step exon recognition is commonly perturbed in cancers

The coordinated AE-IR regulation is observed not only in RON but also transcriptome-wide across human tissues and, remarkably, it is altered in many cancers (Figure 1C and Supplementary Figure S1). Intron retention is known to strongly accumulate in most cancer types compared with other splicing events (9). We confirmed that this result also holds in complex splicing events combining IR and AE, where the change of IR level (median value up to 9%) is in general larger than that of AE (median value < 5%) in each cancer type (Figure 6A).

**Figure 6.**
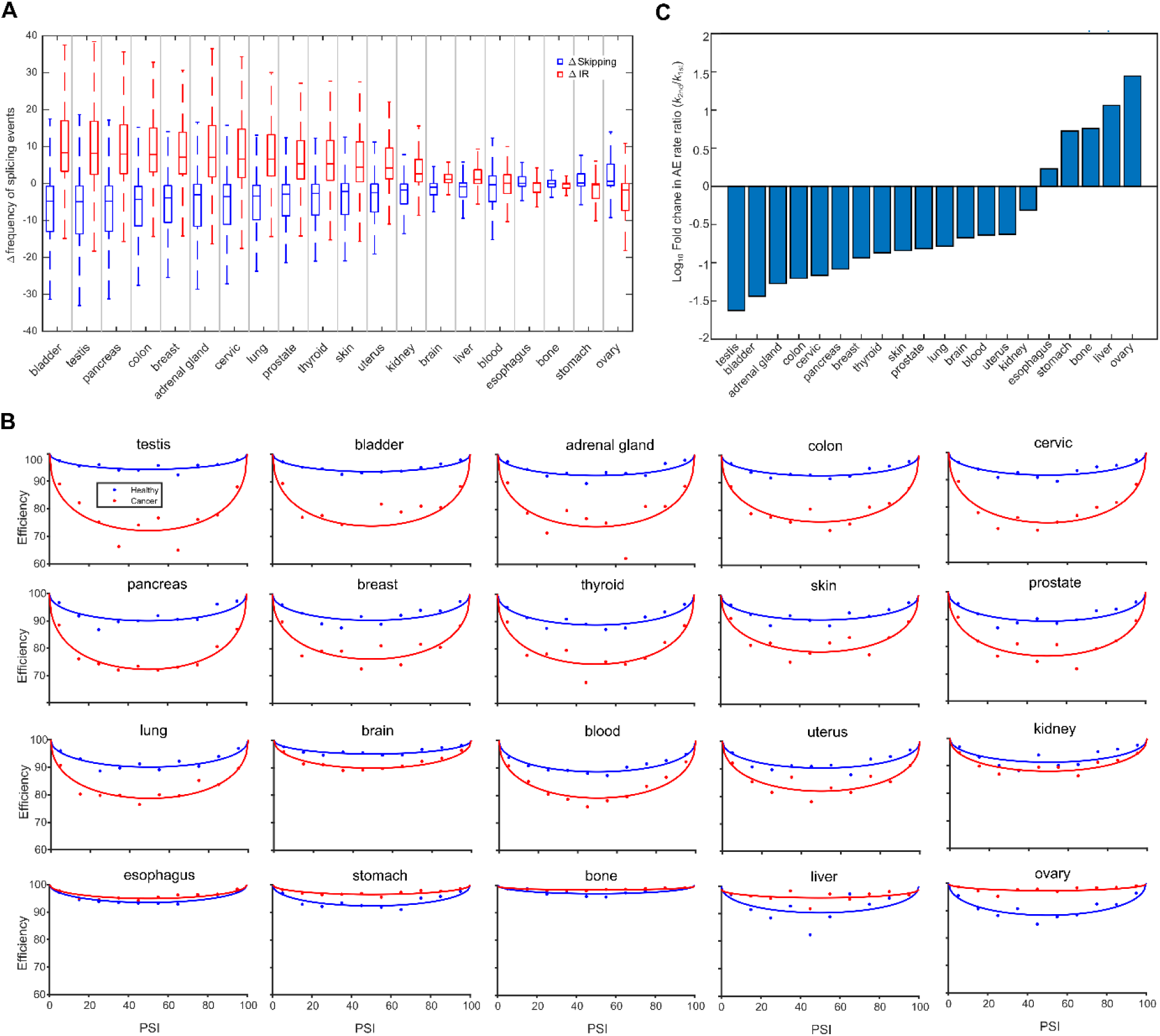
AE definition steps are differentially perturbed in cancers. **(A)** IR is preferentially affected in cancers. The MAJIQ-processed transcriptome-wide RNA-seq data were collected in 20 human tissues (normal tissue data from GTEx, BLUEPRINT, Leucegene and Ranzani et al.; cancer data were from TCGA, TARGET, BEAT-AML and CBTN different healthy tissue sub-brain regions). Splicing events with combined AE and IR decisions were selected for further analyses. Within the combined AE and IR splicing decisions, IR (red) level increases and AE skipping (blue) declines in most cancer types. **(B)** A core two-step AE definition model explains the tissue-dependent efficiency-PSI relation shift in cancers. The splicing efficiency drops at intermediate PSI level in 15 out of 20 tissues (first 3 rows), whereas 5 other tissues showed the opposite effect (the last row). To quantify this tissue-dependent transcriptome-wide efficiency-PSI relation, a simplified two-step model was derived from the full model for RON minigene (Methods). This core model was very well fitted to the data (median efficiencies in PSI bins) in 20 normal (blue) and cancerous (red) tissues. **(C)** AE definition steps were differentially modulated in cancers. The parameter inference from the best fits in (B) showed that the relative rate (or rate ratio) of the two AE definition steps is most significantly altered across all 20 cancer types. 15 out of 20 cancer types showed an accelerated early or/and slowed late AE definition step (a decrease in the ratio between the rates of second and first step), while other 5 cancers showed the opposite (an increase in the rate ratio).

To quantify and explain the transcriptome-wide coordinated AE-IR deregulation in cancers, we analyzed a core model of two-step AE definition simplified from the full model for the RON minigene (Figure 2A). This core model describes the definition of all three exons, which is more generic than the multi-step toy model with constitutive outer exons (Figure 3B). Here, we assumed that all reversible exon definition steps reach equilibrium rapidly and that the two outer exons share the definition rate (Methods).

To identify the regulatory mode(s) shifting the AE-IR coordination in cancers, we fit the core model to the data, using median splicing efficiencies in binned PSI intervals for data from both, normal and cancer tissues (Methods). The core model well captured the efficiency-PSI dependency in all 20 normal and cancer tissues (Figure 6B). The most varied model parameter (from the best fit) between cancer and normal tissues is the ratio of the two AE definition rates, suggesting that the first and second AE definition steps are differentially perturbed in cancers (Supplementary Figure S4). Interestingly, this differential AE modulation is fine-tuned to exhibit distinct IR accumulation patterns in different tissues (Figure 6C). 15 out of 20 tissues showed a slower late or a faster early AE definition step in cancer, which links to a general increase in the amplitude of the efficiency change (with PSI). The remaining 5 tissues showed the opposite effect. Tissues such as in kidney and esophagus, showed small change (< 2 fold) in AE definition steps and thus the AE-IR dependency is unaffected. When the change in rate ratio is mild (2 to 10 folds), the IR (or efficiency) shifts almost uniformly across different AE levels (breast, thyroid, skin, prostate, lung, brain, blood and uterus with increase in IR; stomach and bone with decrease in IR). However, in some tissues the IR preferentially accumulated or declined at intermediate AE inclusion levels, as the two AE definition steps were very distinctly affected in cancers (testis, bladder, adrenal gland, colon, cervix and pancreas with IR increase; liver and ovary with IR decrease).

Taken together, we quantified the AE-dependent IR events as a global trend at the transcriptome-wide level and showed that the perturbation of IR in cancers can be explained by modulation on only a small number of the rate-limiting steps in AE definition, implying a potential direction to search for the common factors responsible for splicing dysregulation in cancers.

## Discussion

Alternative splicing diversifies the human transcriptome by producing different RNA isoforms from the same gene. Traditionally, AE inclusion/skipping has been the focus of splicing research as it is the most common AS event, and the binary AE choice is quantified by a single metric (PSI). On the other hand, intron retention was largely overlooked until recently, where its functional significance was revealed in gene regulation, cellular responses and differentiation (1–3, 6, 7). AE and IR events are controlled by the same molecular machinery, the spliceosome. However, how AE and IR are coordinated is still an open question, even though transcriptome-wide data showed that both events frequently co-occur at the same exons (24, 25). In this work, we showed that AE and retention of its flanking introns are coordinately modulated, where IR accumulates preferentially at intermediate AE inclusion levels. This AE-IR dependency was observed in large-scale mutagenesis library of RON minigenes as well as transcriptome-wide in 20 human tissues. Importantly, our data indicates that altered spliceosome activity does not only change the overall splicing rate (i.e., the sum of exon skipping and inclusion), but that retention is additionally coupled to the skipping/inclusion ratio. To explain this phenomenon, we implemented mechanistic models describing stepwise spliceosome maturation.

We built our models based on the splicing outcomes of the three-exon RON minigene, which leads to the production of five isoforms, AE inclusion and skipping as well as three intron retention isoforms. To study how the spliceosome controls these splicing decisions, we tested two model variants. In the one-step exon definition model, the direct assembly of the early U1-U2 complex cross each exon is the only rate-limiting step of spliceosome regulation. In contrast, the two-step model describes further spliceosome commitment and maturation by incorporating an additional rate-limiting step (Figure 2A). Our analysis showed that the one-step model only allows for independent regulation on AE and IR decisions (Figure 2C). On the other hand, the two-step model by nature enables IR accumulation (or drop in splicing efficiency) at intermediate AE inclusion levels, therefore better accounting for the data of all five RON isoforms (Figure 2D). The two-step model is a minimal representation of the progressive cycle of spliceosome assembly, remodeling and activation. While the spliceosome maturation was often characterized as the behavior of the cross-intron complex, recent studies revealed that the formation of cross-exon spliceosome is highly similar to that of the cross-intron complex, including U1-U2, U4/U6.U5 tri-snRNP assembly and their compositional remodeling (13–15). This supports our model assumption that multiple rate-limiting steps may exist in the exon definition process.

We further provide evidence that multistep exon definition establishes a splicing switch, in which moderate changes in AE recognition are sufficient to induce pronounced transition from complete skipping (PSI = 0%) to complete AE inclusion (PSI = 100%) (Figure 3A). Therefore, multi-step spliceosome maturation provides a kinetic-proofreading-like mechanism that disfavors a splicing decision for exons with intermediate AE recognition, and instead leads to spliceosome rejection, intron retention and potentially non-sense mediated transcript degradation. By exploring a toy model, we showed that the sharpness of the splicing switch increases with the number of proofreading (exon-definition) steps, but at a cost of losing splicing efficiency (Figure 3C and D). This suggests that a small number of rate-limiting steps were used in the spliceosome assembly and maturation to balance the tradeoff between the sensitivity and speed of the splicing switch.

The degree of AE-IR dependency is subject to regulation, as shown by the varied amplitude of splicing efficiency drop in RON minigene upon RBP KDs (Figure 4). Qualitative simulations and systematic model discrimination revealed that the efficiency drop at intermediate PSIs is most efficiently perturbed by a differential modulation on the two steps of AE definition: The drop is most pronounced when the late step is much slower than the early step during spliceosome assembly and maturation, as this leads to more pronounced ‘proofreading’ of the AE. More specifically, it is the ratio of the first and second definition rates rather than their absolute values that determines the amplitude of the drop. Therefore, a strong efficiency decline can result from RBP KD that either speeds up the first step, e.g., SMU1 KD (Figure 5B), or slows down the second step, e.g., SRSF2 KD (Figure 5B). In addition, both SMU1 and SRSF2 were also inferred to regulate the two outer exons, but this type of regulation generally lowers the splicing efficiency across all PSI values. Interestingly, a recent study conducting systematic KDs of ∼ 300 spliceosome components and RBPs reported that KDs of factors involved in the late stage of spliceosome cycle preferentially led to elevated IR accumulation transcriptome wide, while early-stage factor KDs showed the opposite effect (16). This IR regulation directionality agrees with our model predictions above, suggesting that the differential regulation on the rate-limiting steps of spliceosome cycle might be a general mechanism (beyond RON minigene) for coordinating splicing decisions.

Indeed, we observed coordinated AE-IR regulation transcriptome-wide in 20 normal and malignant human tissues and found that the splicing efficiency at intermediate PSIs is altered in most cancers. Interestingly, the majority of the cancer types (15 out of 20) showed a decrease in splicing efficiency, while a small subset (5 out of 20) showed the opposite tendency. This tissue-specific variation of the efficiency decline can be quantitatively explained by a differential modulation of the two AE definition steps in cancer. In the same vein as in the RON minigene, if the late step becomes slower or the early step becomes faster in cancer compared to its normal counterpart, IR will further accumulate at intermediate PSI. With opposite modulations, IR decreases instead, as observed for the minority of the cancers. The detailed molecular mechanisms causing this differential modulation of the two AE recognition steps remain unknown. It is possible that specific factors regulating the steps of AE definition are commonly mutated in many cancer types, such as SF3B1 (33–38). SF3B1 regulates the stepwise U2 assembly and 3’SS commitment till the late stages of spliceosome cycle, and its mutation could strongly disturb the balance between early and late effects. Another possibility is that the relative abundance of different RBPs fails to maintain in cancer due to loss in homeostasis of gene expression (39, 40), and it therefore causes unbalanced controls in distinct splicing steps. In addition, early and late spliceosome states may rely on different energy supplies, where the late stage of spliceosome commitment and remodeling cost more ATPs than the early stage (41–43). In cancer cells, energy supply is generally in high demand for many upregulated cellular processes and may be limited for splicing. Therefore, the energy demanding late step of spliceosome maturation might be more sensitive to the reduced energy supply, resulting preferentially in IR accumulation. Further experiments and analyses would be needed to identify the detailed molecular mechanisms.

In total, we showed how AE and IR decisions are coordinated by the differential modulation of rate-limiting steps during spliceosome assembly and maturation. We expect that the stepwise model could be further extended to include additional types of splicing decisions, such as alternative 3’ and 5’ splice-site choice or mutually exclusive AE, so that the coordination of more complex splicing events can be studied in normal cell physiology as well as in search of novel therapeutics for diseases like cancer.

